# Environmentally-mediated yield effects of vernalization and photoperiod alleles in historic winter wheat trials

**DOI:** 10.64898/2026.09.18.752491

**Authors:** Noah DeWitt, Zachary Winn, Mohammed Guedira, Jeanette Lyerly, Christian Maltecca, Brian Ward, William Stafstrom, Md. Ali Babar, Nonoy Bandillo, Richard Boyles, Stephen Harrison, Kimberly Howell, R. Esten Mason, Ellen Melson, Mohamed Mergoum, Mohsen Mohammadi, J. Paul Murphy, Eric Olson, Jessica Rutkoski, Nicholas Santantonio, Jared H. Smith, Clay Sneller, Mark Sorrels, Russel Sutton, Vijay Tiwari, David A. Van Sanford, Gina Brown-Guedira

## Abstract

In common wheat (*Triticum aestivum* L.), variation at the *Vrn1* and *Ppd1* loci changes plant phenology, resulting in differential adaptation useful to breeders. Because the effects of phenology on grain yield are conditional on environmental factors, the incorporation of markers for *Vrn1* and *Ppd1* alleles into breeding approaches has proven difficult. Historic phenotypic, genotypic, and environmental data can provide insight into the relationship between major phenology alleles and grain yield as mediated by environmental variables. Analyses of eight years of breeding trials (1,038 lines at 219 site-years) determined that weak *vrn1* alleles and *Ppd1* insensitivity alleles have variable effects. These effects change depending on an environment’s winter temperature and latitude, respectively, but the environmentally-driven changes in effect of *vrn1* alleles are much more variable than effects of *Ppd1* alleles. Mediation analyses showed phenologically-dependent environmental variables are one mechanism through which phenological changes created by *vrn1* and *Ppd1* variants alter yield across multiple developmental stages. Environmentally-driven relationships between flowering time and yield were modeled to estimate yield allele effects as a function of effects on phenology and environmental variables. Weak winter alleles at *vrn1* may play a stabilizing role on yield, due to the correlation of winter conditions that drive larger effects with later-season warm temperatures that penalize later maturity. A better mechanistic understanding of how these loci drive environment-specific yield effects may facilitate utilization of markers for these alleles and improve predictive modeling of yield.

## Introduction

Wheat (*Triticum aestivum* L.) production in the eastern United States stretches from the Gulf Coast to the western shore of Lake Huron. Common wheat’s ability to respond to environmental signals facilitates this wide range; an adapted winter wheat genotype uses environmental signals accumulated during the winter and early spring to break dormancy, flower, and then fill grain within an optimal date range that maximizes grain yield. In this way, the genotype determines the environment it experiences at specific developmental stages, and adapted genotypes position themselves to experience favorable environmental conditions during flowering and grain fill. Individual breeding programs face the challenge of selecting plants with phenological profiles suited for their target environments, relying on visual observations of maturity in early generations, and multi-location yield data in later generations. The introduction of genomic selection has extended the prediction of additive merit for multi-location yield to earlier generations, but models average across the environments sampled in their training population. Average additive values collapse year-to-year variability into single values and ignore the underlying biology driving aggregate yield differences. If the environments in the training population are not representative of the breeder’s target population of environments, predictions from these models may actually impede the breeder’s goal of identifying lines with appropriate adaptation.

Better modeling of environment-specific yield effects will necessitate modeling of the mechanisms through which phenology genes generate yield variation. Alleles with yield effects drive differences in the physical structure of plants that may alter their interactions with their surrounding environment. As plants grow within their surroundings, allelic variation in phenology genes alters plant response to environmental conditions, which in turn alters the environment they experience at their next developmental stage (Lewontin, 1957; Allard and Bradshaw, 1964). In winter wheat, variation in heading date is used to track the general differences in plant phenology produced by these interactions. Heading date occurs a few days prior to flowering, associated with the emergence of the wheat spike, and is correlated with various measures of plant developmental rate, including dormancy release and the start of grain filling. Differences in developmental rate expose plants to different environmental conditions during comparable reproductive stages. Earlier maturity tends to reduce grain size by shortening the grain fill period during which carbohydrates are allocated to developing grain (Sofield et al., 1977). Moreover, early heading date may also reduce yield by exposing developing spikes to below-freezing temperatures that render florets infertile (Paulsen and Heyne, 1983). In contrast, heading too late risks reducing yield by exposing plants to heat or drought stress during grain fill. Post-heading abiotic stress can reduce grain size by preventing the movement of carbohydrates into grain, increasing root and shoot temperatures, decreasing sucrose levels, and deactivating the starch synthase enzyme that assimilates carbon, which is particularly heat-sensitive in wheat (Bhullar and Jenner, 1985; Hawker and Jenner, 1993; Guedira and Paulsen, 2002). Differences in plant phenology genes influence their responses to their environments, driving differences in experienced environmental conditions across developmental stages, which can then drive differences in grain yield.

In wheat, several alleles affecting core genes in the flowering time pathway have been used by breeders to alter heading date. The transition of fall-planted wheats from dormancy to reproductive growth results from the expression of the ortholog of the *FLOWERING LOCUS T (FT)* protein (*VRN3* in wheat) and its movement through the phloem from the leaves to the shoot apical meristem (Yan et al., 2006; Gauley and Boden, 2019). Transcription of *Vrn3* is initiated through activation of a set of upstream genes which integrate signals from night length (the photoperiod pathway), accumulated chilling hours (the vernalization pathway), and the age of the plant in units of thermal time (the earliness per-se pathway). The major heading date genes of wheat involve alleles at the A, B, and D homeologues of the major integrator of vernalization signal, *Vernalization1* (*Vrn1*) (Yan et al., 2003, 2004; Fu et al., 2005; Li et al., 2013), and the major integrator of photoperiod signal, *Photopeiord1* (*Ppd1*) (Turner et al., 2005; Beales et al., 2007; Nishida et al., 2013; Guedira et al., 2016), both upstream of *Vrn3* in the wheat flowering time pathway. Dominant alleles conferring spring growth habit (zero chilling hour requirement) constitutively up-regulate the *Vrn1* loci on the chromosome 5 homeologues (Yan et al., 2004; Fu et al., 2005; Li et al., 2013). Alternative recessive alleles of *Vrn1* in winter wheat (*vrn1* alleles) reduce the degree of vernalization sensitivity and promote earlier flowering, especially in warmer climates (Fu et al., 2005; Díaz et al., 2012; Guedira et al., 2014, 2016; Kippes et al., 2018). Variation in copy number and SNPs in exon4 and exon7, as well as regulatory sites of the *VrnA1* first intron, have been associated with differences in vernlalization requirements in winter wheat (Díaz et al., 2012; Li et al., 2013; Kippes et al., 2018). Likewise, a 36 bp deletion in *VrnB1* intron one was associated with reduced vernalization requirement (Guedira et al., 2014, 2016). A wheat with one or more vernalization alleles for reduced sensitivity may be termed a “weak”winter wheat, as opposed to a “strong”winter wheat with a larger chilling hour requirement.

Photoperiod response is also a critical factor affecting adaptation and flowering in wheat. In photoperiod sensitive wheat, flowering is accelerated by exposure to short nights (long days).*Ppd1* genes on all three chromosome 2 homeologues (*Ppd-A1, Ppd-B1, Ppd-D1*) condition wheat’s response to night lengths (Beales et al., 2007). Misexpression of these genes caused by large deletions upstream of the *Ppd-D1* and *Ppd-B1* loci lead to insensitivity (Beales et al., 2007; Nishida et al., 2013). Any insensitivity allele (referred to as an “a”allele) at *PpdA1, PpdB1*, or *PpdD1* is sufficient to induce a photoperiod insensitive habit, removing the requirement of shorter nights to trigger the transition to reproductive growth, and then have different quantitative effects on heading date in an already photoperiod insensitive background (Law et al., 1978; Law and Worland, 1997). In the diverse environments of the Eastern winter wheat growing region, multiple alleles at *Ppd1* for photoperiod insensitivity and *vrn1* for variable vernalization (chilling hour) requirement have been used by breeders to adjust wheat phenology. Although genotypes are routinely screened with available DNA markers for those alleles, the use of this genotyping data for breeding is not straightforward because the complicated relationship between these alleles and grain yield is not well characterized.

Understanding the effects of vernalization and photoperiod loci on local yield performance requires examination of the way environmental variables mediate environment-specific yield effects of heading date variation. To assess these effects, a large number of lines with differing allelic states must be exposed to different environmental conditions over many years and locations. Data on many previous years of cooperative breeding trials in the Eastern United States, aggregated into a combined historic yield dataset, provides such a resource. The use of historic yield trial data is imperfect due to error, lack of balance, and population structure; however, these historical datasets span a vast number of environments over an extended period of time, making them at present the only possible datasets useful to begin addressing these questions (Boyles et al., 2019; DeWitt et al., 2023). In the present study, we perform an exploratory analyses of historic winter wheat trials to better understand the interplay of photoperiod and vernalization allelic states and grain yield as mediated by environmental variables. These results serve as a framework for modeling how individual genes can generate genotype-by-environment (GxE) interactions through altering the interactions between plants and their environments.

## Materials and Methods

### Plant Material and Phenotypic Data

Historic data was curated from 2012-2020 for a series of 1,038 lines tested in a series of cooperative Soft Red Winter Wheat (SRWW) breeding nurseries in the Eastern United States that were genotyped with gene-based markers for *vrn1* and *Ppd1* alleles (Supplemental Table S1). Breeding programs in the Southern region principally contribute to the Southern UNiversities Wheat Nursery (SUNWheat) and the Gulf Atlantic Wheat Nursery (GAWN). Lines tested in the SUNWheat nursery were contributed by members of the SUNGrains cooperative (University of Arkansas, Clemson University, University of Florida, University of Georgia, Louisiana State University, North Carolina State University, and Texas A&M University). The GAWN consisted of lines from the same programs, with the addition of germplasm from the Virginia Polytechnic Institute and State University small grains breeding program. The Mason-Dixon nursery consists of lines from the Virginia Polytechnic Institute and State University, University of Kentucky, USDA-ARS Raleigh, and University of Maryland breeding programs. Entries in the Five-state preliminary and Five-state advanced were contributed by breeding programs at the University of Kentucky, University of Illinois, The Ohio State University, Michigan State University, and Cornell University. For these breeding nurseries, genotyping for both major genes and genomewide markers was performed by the USDA-ARS Eastern Regional Small Grains Genotyping Lab in Raleigh, North Carolina. Breeding programs frequently use lines from other Eastern breeding programs in crossing, so that breeding programs do not represent distinct populations.

In addition to the collaborative breeding nurseries, data from two USDA-ARS Uniform nurseries were used: the Uniform Eastern Soft Red Winter Wheat Nursery (UESRWWN) and the Uniform Southern Soft Red Winter Wheat Nursery (USSRWWN). In both cases, one to four lines from each breeding program that were previously screened in the aforementioned collaborative nurseries are advanced to a Uniform Nursery and evaluated across many sites within a given year. Northern programs submit entries to the UESRWWN, and southern programs to the USSRWN, with some programs contributing to both. These nurseries generally feature fewer lines (28-46) and more environments (13-27) than the breeding nurseries. Genotyping data for major genes is also available for lines in the Uniform nurseries, as well as genome-wide data for entries that were previously evaluated in a collaborative nursery.

These nurseries together capture a set of genotypes and environmental conditions broadly representative of Eastern winter wheat germplasm and target environments. Nurseries were unbalanced across years but balanced within years, consisting of a small set of shared checks planted across years and a larger set of advanced stage lines planted in all environments in a single year. While some advanced lines were trialed in the nurseries in multiple years, typically a line was only screened in a given nursery in a single year, with some entries moving from the earlier-stage breeding nurseries to later stage and uniform nurseries.

Yield trials consisted of plots that were at minimum 1.3 m wide and 3.1 m long and managed following local recommendations. Yield was taken as bushels per acre, assuming a constant test weight of 60 lb bu^*−*1^ and adjusting for moisture content as appropriate, and converted into kg ha^*−*^1 on the rate of 67.25 kg ha^*−*^1 per bushel per acre. Heading date was reported for plots as the day of the year past January 1 at which approximately half of the heads of the primary tillers in a plot had fully emerged from the boot. Entry means for each genotype in each trial were reported by collaborators following individual analyses. Data from field sites that were damaged or with major planting or harvest errors was not reported. After removing lines without yield or heading date data and other filtering for data quality, the final data set consisted of 13,002 observations of 1,008 distinct genotypes.

### Climate Data

Latitude and longitude coordinates for each field location were aggregated based on information from collaborators and observations of wheat trials in Google Maps (https://maps.google.com) imagery. Daily meteorological data (relative humidity, specific humidity, mean temperature, max temperature, minimum temperature, short wave solar radiation, long wave solar radiation, atmospheric pressure, precipitation, and mean wind speed) for each latitude-longitude pair were downloaded from the NASA POWER data set using the *nasapower* package in R (R Core Team, 2021; Sparks, 2018). In this study, we refer to two types of environmental variables were calculated from publicly-available weather data: phenology independent (PI) variables, which are associated with a site-year and are consistent for every plot in a given trial, and phenology dependent (PD) variables, which are calculated based on the relative heading date of individual plots, and are different for each individual plots within a site-year based on crop modeling of developmental periods. Developmental periods were calculated for the trial as a whole and for individual heading dates within a trial based on re-implemented formulas used by the APSIM crop modeling wheat module (Zheng et al., 2015).

Mean daily temperature *T*_*avg*_ was taken as the average of daily maximum and minimum temperatures. Daily change in vernalization accumulation, Δ*V*_*ij*_, was taken following the CERES approach as the minimum of either 1.4 *−* 0.0778*T*_*avg*_ or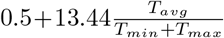, which accounts for extremes in minimum and maximum daily temperature, when both *T*_*max*_ *<* 30°*C* and *T*_*min*_ *<* 15°*C*. Daily change in thermal time Δ*R* was calculated as daily average temperature *T*_*avg*_ when *T*_*avg*_ *>* 0°*C* and *T*_*avg*_ *≤* 26°*C*, and as 3.25(34 *− T*_*avg*_) when *T*_*avg*_ *>* 26°*C* and *T*_*avg*_ *≤* 34°*C*. Freeze damage was calculated as the degrees Celsius beneath freezing up until *−* 12°*C*, at which point no additional freeze damage is expected. Within each site-year, mean and total meteorological variables were calculated across the winter season prior to winter dormancy release. Approximate date of winter dormancy release for wheat in a given site-year, used to determine the approximate time period across which phenologically independent values were summed, was calculated using default values from the APSIM (Agricultural Production Systems sIMulator; https://www.apsim.info/) wheat module. Briefly, after the first day of the year change in adjusted thermal time *R*^*′*^ for each day *i* within each site-year 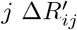 was calculated as:

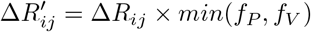

Such that either insufficient day length or accumulated vernalization (represented by photoperiod adjustment parameter *f*_*P*_ or vernalization adjustment parameter *f*_*V*_) delays growth. Daily photoperiod adjustment within each site year reduces changes in daily thermal time at values of day length *L*_*i*_ lower than 20 hours (*f*_*Pi*_ = 1 *−* 0.006(20 *− L*_*i*_)^2^). Daily vernalization adjustment with each site year reduces changes in daily thermal time when accumulated vernalization *V*_*i*_ is less than 50 (*f*_*Vi*_ = 1 *−* 0.02(50 *− V*_*i*_) with *f*_*Vi*_ constrained to *f*_*Vi*_ *>* 0). Approximate date of winter dormancy release for each site-year is calculated according to APSIM defaults as the first Julian day of the year on which 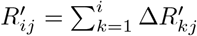 was greater than 400°Cd.

Dates of developmental stages prior to and after heading were estimated for each plot within each site year using plots’ heading date values and working backwards and forwards until standard APSIM thermal time requirements are reached (from floral initiation to flowering, 555 °Cd;from flowering to grain fill initiation, 120 °Cd; from grain fill initiation to termination, 500 °Cd). Developmental stage-specific meteorological data for the phases floral development, flowering, and grain fill were calculated by summing, averaging, or calculating the maximum value for meteorological variables within each developmental stage.

As soil moisture has a more direct relationship to grain yield than precipitation totals, a simplified soil moisture model was constructed for each site. Briefly, data on soil composition for each field site was obtained using the *soilDB()* package in R (Dylan Beaudette et al., 2021). Sand, silt, and clay fraction along with soil bulk density were used to calculate site-specific percent soil moisture values for field and wilting capacity using the implemented Rosetta pedotransfer model Zhang and Schaap (2017). For each day, meteorological values and latitude information were used to estimate daily evapotranspiration for a short grass crop according to the Penman-Monteith model (Allen et al., 1998).

### Genotypic Data Collection and Analysis

Each year, DNA of nursery entries was isolated from a bulk of tissue from a six plants using the sbeadex plant maxi kit (LGC Biosearch Technologies, UK). DNA was normalized to 20 ng/ml for downstream marker analyses. Entries were evaluated with a series of Kompetitive allele specific PCR (KASP) assays targetting polymorphisms at the *Vrn1* and *Ppd1* loci (Supplemental Table S1). KASP markers for *vrnA1* target SNP in *VrnA1* exon 4 and exon 7 differentiating haplotypes that are also associated with copy number differences, where strong winter types at the locus have three or more copies of the*vrn1* winter allele and weaker winter types have one or two copies (Díaz et al., 2012; Kippes et al., 2018; Jiao et al., 2025). Lines were also assessed with an assay targeting a 36 bp deletion in the first intron of the winter allele of *vrnB1* loci associated with reduced vernalization duration requirement (Guedira et al., 2014, 2016). To assess photoperiod insensitivity, assays targeted large deletions upstream of the *PpdD1* and *PpdA1* loci (Beales et al., 2007; Nishida et al., 2013). For the *PpdB1* locus, assays targeting intercopy junctions were used to identify lines having reduced photoperiod sensitivity due to the presence of more than one *PpdB1* copy (Díaz et al., 2012). Sequences of KASP primers are presented in Supplemental Table 1. PCR protocols followed manufacturer’s instructions, which consisted of a reaction volume of 4.0 *µ*L comprised of 2 *µ*L 2x KASPar reaction mix, 0.05 *µ*L 72x assay mix, and 2 *µ*L of template DNA (10 ng *µ*L^*−1*^). Endpoint fluorescence was analyzed using the KlusterCaller genotyping software (LGC Biosearch Technologies, UK).

**Table 1.** Frequencies of early-flowering alleles in phenotyped testing set. Total number of lines assayed for phenotype and genotype at all five loci in all nurseries (northern nurseries: GAWN, SUNWheat, USSRWWN, MidAtlantic nurseries: Mason-Dixon, ABB Mid-Atlantic, Northern Nurseries: Five State Pre and Adv, UESRWWN). Allele frequencies of the allele associated with early heading is given for each locus and nursery, along with across-homeologue, multi-locus haplotypes.

| Nursery | *n* | *Ppd1* insensitive ("a") |  |  | *n* of *Ppd* "a" copies |  |  |  | weak *vrn1* |  | *n* of *vrn1* |  |
| --- | --- | --- | --- | --- | --- | --- | --- | --- | --- | --- | --- | --- |
|  |  | *A* | *B* | *D* | 0 | 1 | 2 | 3 | *A* | *B* | weak | strong |
| GAWN | 199 | 0.46 | 0.15 | 0.68 | 1 | 108 | 61 | 15 | 0.21 | 0.19 | 78 | 121 |
| SUNWheat | 335 | 0.42 | 0.12 | 0.68 | 1 | 164 | 103 | 27 | 0.27 | 0.30 | 185 | 150 |
| USSRWWN | 174 | 0.46 | 0.09 | 0.63 | 0 | 110 | 41 | 9 | 0.19 | 0.18 | 68 | 106 |
| Mason-Dixon | 128 | 0.50 | 0.12 | 0.78 | 1 | 62 | 48 | 3 | 0.17 | 0.04 | 30 | 98 |
| ABB Mid-Atlantic | 67 | 0.60 | 0.16 | 0.81 | 1 | 18 | 37 | 6 | 0.10 | 0.10 | 16 | 51 |
| Five State Pre | 145 | 0.73 | 0.12 | 0.57 | 4 | 68 | 54 | 5 | 0.01 | 0.00 | 1 | 144 |
| Five State Adv | 144 | 0.69 | 0.15 | 0.52 | 3 | 77 | 45 | 4 | 0.01 | 0.02 | 5 | 139 |
| UESRWWN | 164 | 0.57 | 0.13 | 0.62 | 0 | 96 | 49 | 4 | 0.07 | 0.02 | 15 | 149 |

Genome-wide markers were obtained for lines through reduced-representation genotyping by sequencing (GBS), using the *PstI* and *MstI* enzymes as described in Poland et al. (2012). Discovery of single-nucleotide polymorphisms (SNPs) was performed through alignment of sequenced short reads to the International Wheat Genome Sequencing Consortium Chinese Spring RefSeq assembly v1.0 (https://wheat-urgi.versailles.inra.fr/Seq-Repository/Assemblies) through the Burrows-Wheeler algorithm (Li et al., 2009) as implemented in the SNP-calling software Tassel (Glaubitz et al., 2014). Sequence data for SNP discovery included all lines in the study along with additional SRWW lines to identify and retain variants with a low minor allele frequency. Variants identified using this discovery set were filtered based on expectations for SNPs in a population of largely inbred lines – SNPs were retained at MAF ≥ 5%, heterozygosity ≤ 10%, and missing data ≤ 50%. Imputation of missing SNPs was accomplished using Beagle V4.2 (check) (Browning and Browning, 2016), followed by a second filtering step using the same parameters. A production run of Tassel was then run for all lines in the data set. Markers were thinned based on LD using the R package *gaston* for *r*^2^ ≤ 0.80 (Dandine-Roulland and Perdry, 2018).

Pairwise realized relationships between lines were estimated from markers as relationship matrix **G. G** was computed in R according to the Van Raden method as 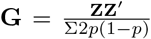 (VanRaden, 2008), where *p* represents a vector of allele frequencies of all SNP in the (-1, 0, 1) coded marker matrix **M**, and **Z** = **M** - **P**, where vector *p* is duplicated according to the rows in **M** to generate matrix **P** of the same dimensions.

### Pedigree Data Collection

Pedigree data for all lines was aggregated from historic entry lists for each year and trial. Pedigree strings in Purdy format were processed using R and converted into a three-column pedigree format, with additional entries added for for intermediate individuals implied by three-way or backcrosses. The three-column format pedigree file was manually cleaned to remove special characters and to strip out additional information (typically, synonym information relating experimental names to variety names or gene information). Sub-string matching was used to generate candidate synonyms between line names found in pedigrees, and candidate synonyms were manually curated and added to the synonym information obtained from pedigree strings to generate a full set of synonyms. Pedigree information on lines in the phenotyped set of trials was combined with pedigrees from the wheatpedigree.net database to generate deep pedigree information. The total pedigree set consisted of 7,308 distinct lines. An additive numerator relationship matrix **A** was estimated from the three-column pedigree set using the *A*.*matrix()* function from the R package *AGHmatrix* (Amadeu et al., 2016).

A majority of lines for which pedigree data was available (5,325 of 7,308) did not have genome-wide SNP data. To include them in the analysis, a single relationship matrix was created for both the genotyped lines represented in both **G** and **A** and the ungenotyped lines represented in only **A**. This blended **H** matrix was calculated in R by sub-dividing **A** into matrices representing relationships among genotyped lines (**A**_11_), ungenotyped lines (**A**_22_), and the relationships between geno-typed and ungenotyped lines (**A**_12_ and **A**_21_). **H** was calculated as:

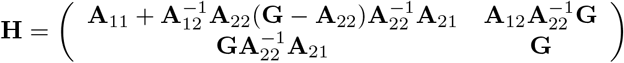

Where the genotyped subset **A**_22_ is replaced by **G**, and adjust estimates of kinship in the remainder of **A** based on the deviation of realized kinship estimates in **G** from estimates in **A**. Principal components of the **H** matrix used to control for population structure were generated with the *prcomp()* function in R.

### Allele Effects for Heading Date Genes

Effects of major *Vrn1* and *Ppd1* alleles on heading date were computed by merging pooled KASP calls for various earliness alleles at the same loci to generate combined, across-homeologue *Vrn1* and *Ppd1* calls. For the two vernalization loci (*vrnA1* and *vrnB1*), lines were grouped into classes based on their alleles at both loci (strong vernalization, *vrnA1* weak, *vrnB1* weak, *vrnA1 + vrnB1* weak). This approach was not taken for the photoperiod loci due to observed disequilibrium between homeologous loci, such that there were almost no true photoperiod sensitive lines in the data set. Across the three homeologous loci, a combined dosage of photoperiod insensitivity alleles was taken; allele dosage of photoperiod insensitivity alleles was measured as the total number of photoperiod insensitive alleles per line (*e*.*g*., 2 for homozygous at one locus, 4 for homozygous at two loci, etc.).

To test the effects of vernalization class and photoperiod insensitivity allele dosage, a mixed linear model was fit using the historic phenotype and marker data. The first six principal components of **H** were included as **Q** and fit as a separate fixed effect for each individual principal components vector *q*:

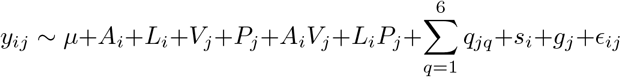

where *y*_*ij*_ is a vector of entry means of genotypes *j* in each site-year *i, A*_*i*_ and *L*_*i*_ represent the accumulated vernalization and latitude of each site-year *i, V*_*j*_ and *P*_*j*_ represent the vernalization class and photoperiod allele dosage of genotype *j, s* represents the IID random effect of site-year *i* across genotypes, *g* represents the IID random effect of genotype *i* across site-years, and *ϵ*_*ij*_ represents the residual for site-year *i* and genotype *j*.

After sufficient vernalization has been accumulated, additional vernalization is not expected to hasten flowering (Flood and Halloran, 1984). In order to better model this relationship, the above model was iteratively rerun, differing only in that all vernalization values above some threshold *t* were set to *t*. The optimal value of *t* (representing the threshold at which additional vernalization no longer hastens heading in our environments and populations) was learned via one-dimensional gradient descent starting from an initial low value of *t* = 20 (vernalization accumulation *V* in calculated ° C) and moving towards a minimum AIC. This optimum value determined via a minimum AIC is equivalent to that determined from a maximum log likelihood. The value of *t* associated with the lowest-AIC model was taken as the accumulated vernalization threshold, and coefficients from that model were used for subsequent analyses of allele effects.

### Variable Selection and Mediation Analysis

A Bayesian lasso approach was used to identify environmental variables driving correlations between heading date and yield in different environments. Lasso differs from ridge regression by assuming a double-exponential distribution rather than a normal distribution, forcing many small-magnitude regressor coefficients to exactly zero (Tibshirani, 1996). Bayesian estimation of lasso regression coefficients generate distributions of coefficients for each feature, allowing for selection of important features by excluding those routinely estimated as having no effect. Coefficients **b** were estimated for **y** *∼* **Xb**+**e**, where **y** is the vector of entry means for yield and **X** represents the matrix of all calculated phenologically dependent variables for the three developmental stages. Solutions were estimated using Gibbs sampling as implemented in the *blasso()* function in R package *monomvn*, and the inclusion probability of weather features was used to select relevant weather variables if they were set to zero in less than 1% of 1000 iterations (corresponding to *α* = 0.01) (Gramacy and Pantaleo, 2010).

To understand how within-location heading date differences have effects on within-location yield as mediated by environmental conditions, a mediation analysis model was fit using relative heading date, relative yield, and phenologically dependent variables across all locations with collected heading date. Means of relative heading date and yield for each entry were calculated by subtracting the mean heading date and yield of each site from the entry means of heading date and yield. The model was fit in the R package *lavaan* as a series of joint regressions of relative heading date on the previously selected photoperiod-dependent variables alongside a multiple regression of those variables on relative yield. The first six principal components of the **H** matrix were included in this regression to control for population structure **Q** as above, so that for each individual PD variable:

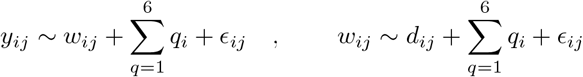

Where *y*_*ij*_ represents the relative yield of line *i* in environment *j, w*_*ij*_ represents the value of an individual PD variable *k* from **W** for line *i* in environment *j*, and *d*_*ij*_ represents the relative heading date of line *i* in environment *j*. Both regression were fit for all photoperiod-dependent variables in **W** simultaneously in *lavaan*. All variables were scaled and centered prior to fitting of the mediation analysis model.

### Trial-Specific Yield Effects of *Vrn1* and *Ppd1* Alleles

A Bayesian lasso model was used to fit a polynomial heading date model with weather-by-heading date deviations to predict the relationship between heading date and yield within each site-year based on environmental variables (Gramacy and Pantaleo, 2010). Site-year-adjusted relative values were used for both yield and heading date as the relationship of interest is the within-site heading date and yield relationship. A numeric matrix **W** was constructed consisting of the subset of *n* phenologically dependent variables identified as important by the previous Bayesian lasso regression, scaled so that each PD vector *w* had a mean of zero and a standard deviation of one. A polynomial trend was fit to accommodate the possibility of an optimal heading date for a given location. An overall trend for heading date effects and interactions with photoperiod-dependent variables in **W** were fit as:

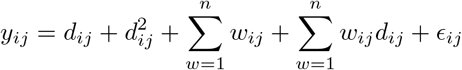

For relative yield *y* and relative heading date *d* for each individual genotype *i* and environment *j*. The model coefficients were multiplied by the observed heading dates and associated photoperiod-dependent variables found within each site-year to predict the yield associated with each and fit a site-year-specific heading date to yield curve. To estimate the yield effects of the heading date alleles as a function of each curve, heading date effects for weak vernalization alleles and number of photoperiod insensitivity allele were calculated for each trial using the previously fit model of allele effects based on accumulated vernalization and latitude. Yield effects for the heading date alleles were estimated by calculating the yield effect associated with their estimated heading date effects centered on the mean trial heading date.

## Results

### Population Structure and Patterns of Major Gene Variation in SRWW

Data was included for individuals evaluated between 2012 and 2020 as part of collaborative evaluation of nurseries targeting the northern part of the US eastern soft wheat growing region (Uniform Eastern, Big Five Advanced, Big Five Preliminary), the Mid-Atlantic region (ABB Mid-Atlantic, Mason-Dixon), and the southern portion of the region (Uniform Southern, Gulf Atlantic Wheat Nursery, and SunWheat). Although the breeding programs that contribute to these nurseries varies, extensive crossing between programs has reduced population structure in the eastern US SRWW region, especially in the Mid Atlantic and South. While analysis of the **H** matrix indicated that substantial population structure exists between lines tested in the Northern and Southern nurseries, ample overlap exists between them, and lines from the Mid-Atlantic nurseries overlap with both (Fig. 2). Individuals tested across multiple nursery groups facilitated estimation of site-year effects.

**Figure 1.**
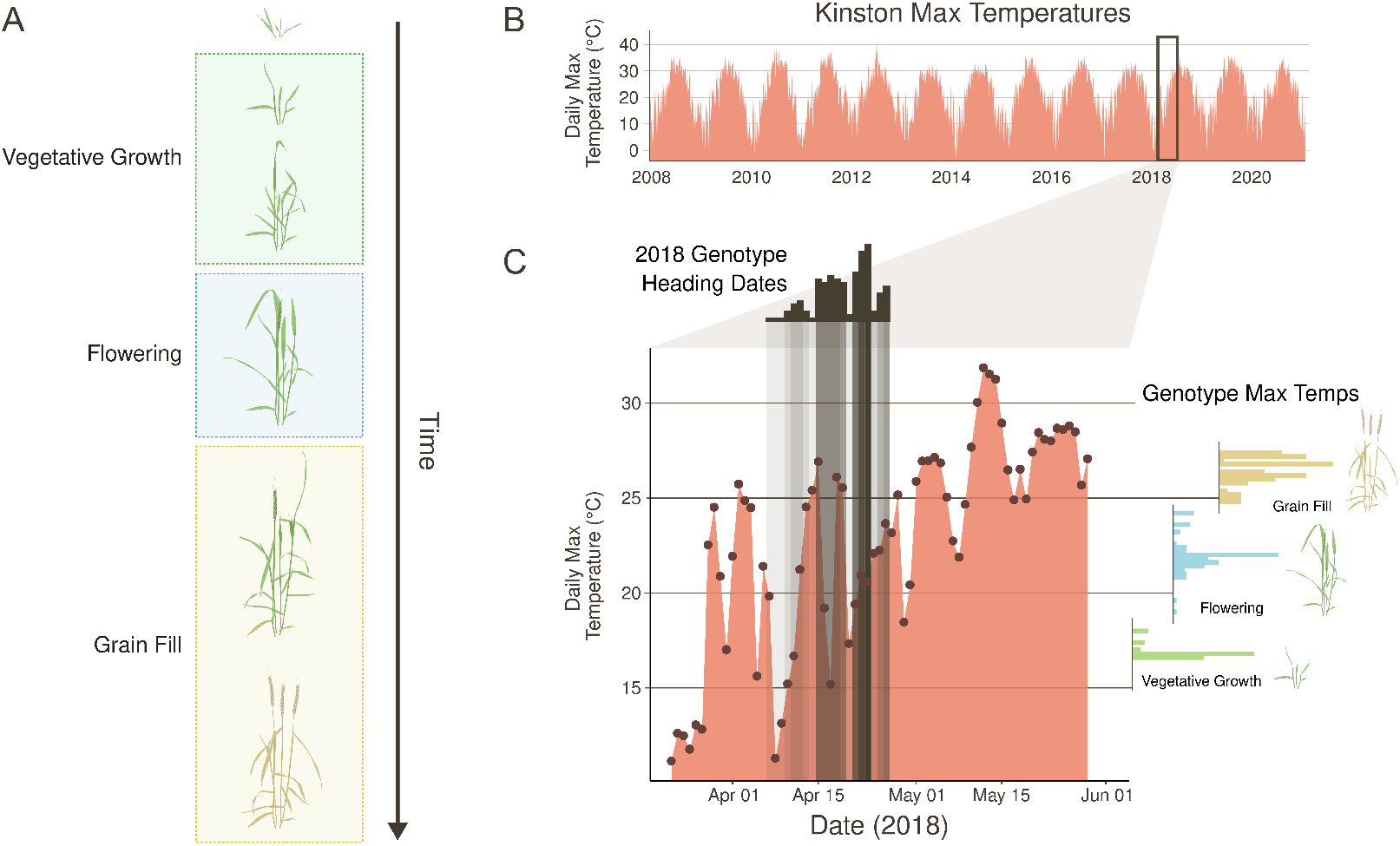
Wheat developmental stages used to calculate per-entry phenologically dependent variables. A - Developmental stage groupings used to summarize variables, defined as before or after heading. B - An example environmental variable (Daily Mean Temperature) over time at an example location (Kinston, NC). C - Different heading dates expose plants to different environmental conditions at different life stages. Earlier headed lines experience different average mean daily temperatures across a range of dates associated with the different developmental stages than later headed lines. Histograms associated with a phenologically-dependent environmental variable calculated from daily max temperature are shown on the right, and correspond to the same y-axis.

**Figure 2.**
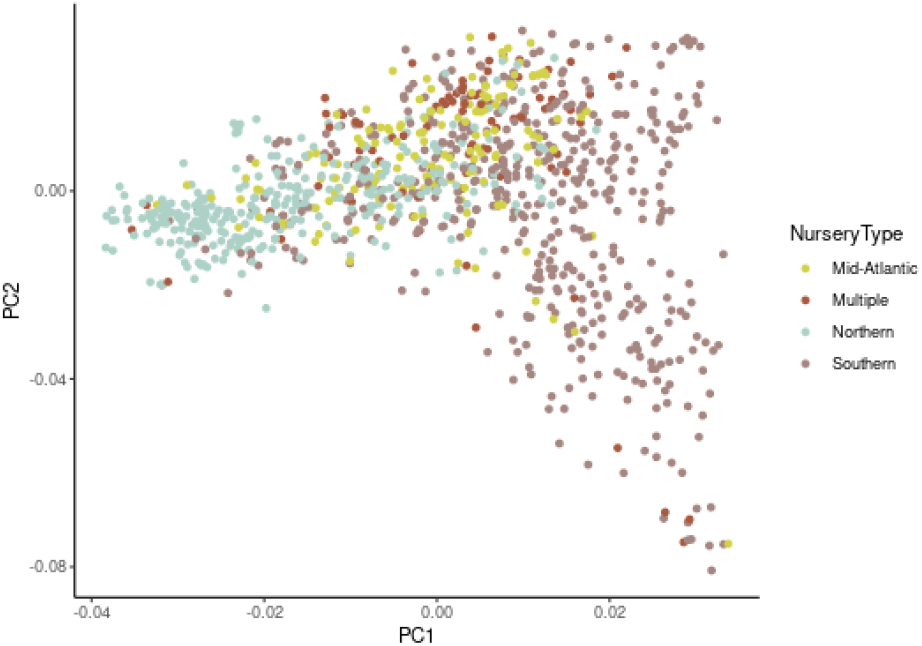
Population Structure in Tested Individuals. First two principal components of **H** matrix showing population structure between individuals in Northern (Uniform Eastern, Big Five Advanced, Big Five Preliminary), Mid-Atlantic (ABB Mid-Atlantic, Mason-Dixon), and Southern Nurseries (Uniform Southern, Gulf Atlantic Wheat Nursery, and SunWheat). Many individuals are found in multiple nurseries, facilitating estimation of site-year effects.

Photoperiod and vernalization alleles were coded by their effect on plant phenology (weak vs strong vernalization requirement, photoperiod sensitive “b”alleles vs insensitive “a”alleles) and aggregated by nursery over the 8-year historic period (Table 1). Allele frequencies for all assayed genotypes, including those with only some loci assayed, are given in Supplemental Table S2. Variation was observed at the three *Ppd-1* loci in all nurseries. Patterns of variation varied by region; comparing the northern nurseries to the southern nurseries, the *Ppd-A1a*.*1* allele was most frequent in northern and Mid-Atlantic lines, particularly entries in the Five State nurseries, but was also common in the southern nurseries. Although this allele was present in more than half of lines assayed in this study, Nishida et al. (2013) found that it was rare in global wheat germplasm, but was described in the historic U.S. eastern soft wheat founding line “Purplestraw”. The potent *Ppd-D1a* insensitivity allele, introduced into U.S. germplasm in the 1960s, was present in 63% to 81% of lines in the southern and Mid-Atlantic nurseries, and at a lower but also majority frequency in the more northern nurseries. Alleles at the *Ppd-B1* locus with insensitivity were substantially rarer, but found in all nurseries at similar rates of 9% to 16%.

In contrast, the frequency of the weak vernalizaiton alleles differed between northern and southern nurseries. The weak winter alleles of *vrn-B1* was present in 0 to 10% of lines from the northern and Mid-Atlantic nurseries, but found in 18% to 30% of lines in the southern nurseries, with *vrn-A1* demonstrating a similar pattern (1% to 17% in the north and Mid-Atlantic vs 19% to 27% in the southern). Similarly, the frequency of vernalization insensitivity at either locus was much higher in the southern nurseries than the northern, constituting a majority of lines in the early-generation SUNWheat, while being found at rates of 10% or lower in the northern nurseries.

Given the high frequency of each individual photoperiod allele, the presence of photoperiod sensitive individuals – lines without any three of the insensitivity alleles – is almost zero (Table 1). The high frequency of individual sensitive alleles and the near-absence of lines with all three alleles for sensitivity suggests a selection against a substantially later habit, regardless of nursery latitude. Testing each allele as a discrete effect is prevented by gametic disequilibrium between loci, likely driven by both this selection and population structure; as an example, there are approximately equal number of photoperiod sensitive and insensitive lines at the *PpdD1* locus within all lines that are *PpdA1* insensitive; within lines that are instead *PpdA1* sensitive, there are almost no *PpdD1* sensitive alleles. This prevents proper estimation of a *PpdA1* effect.

### Heading Date Effects Vary with Cumulative Vernalization and Latitude

Allelic disequilibrium between the three *Ppd1* homeologues complicates allele effect estimation. Further complicating estimation is the known epistatic effects within each group; a single photoperiod insensitivity allele is sufficient to induce a qualitative change from wild-type photoperiod sensitivity to a photoperiod insensitive response, while additional insensitivty alleles at the other loci are associated with a quantitative decrease in heading date. Likewise, a single insensitivty allele at either of the *vrn1* loci is sufficient to induce a qualitative change in the plant response from possessing a strong vernalization requirement to a weak winter type.

Because of the pattern of segregation of the two homeologous allele sets in Eastern SRWW breeding populations, alternative encodings were used to estimate the effects of the *vrn1* alleles and *Ppd1* alleles. In the case of vernalization, these encodings were used to classify lines into four classes: vernalization sensitive, *vrnA1* insensitive, *vrnB1* insensitive, and *vrnA1 + vrnB1* insensitive. These encodings allow for the comparison of different types of insensitivity to a sensitivity baseline. In the case of photoperiod, effects were estimated excluding the small number of truly photoperiod sensitive genotypes (photoperiod sensitive alleles at all three loci). Instead, *Ppd1* genotypes were encoded by the total genotype dosage of insensitivty alleles across all three loci, ranging from the case of a line with a single genotype (2) to a genotype homozygous for insensitivity alleles for all three loci (6), allowing for the calculation of the marginal effect of an additional *Ppd1* insensitive allele.

To understand the effect of environmental conditions on the magnitude of the heading date allele effects, interactions between the vernalization allele classes and a site-year’s accumulated winter vernalization were tested, along with the interaction between additional photoperiod insensitivity alleles and latitude as a proxy for night length. While the data set includes 445 total site-years, individual genotypes were assayed for all relevant KASP markers only after 2012, leaving 219 total site-years in the filtered set used to analyze *vrn1* and *Ppd1* allele effects specifically. In general, locations further north received greater amounts of average accumulated vernalization (Fig. 3). Within each location, however, there was substantial differences in vernalization, depending on the year. Increasing vernalization did not scale exactly with increasing latitude, however, due to the modeling assumptions of APSIM that vernalization is optimally accumulated in the temperature range just above freezing, and decreases at progressively colder temperatures. Some further north locations, especially Nairn, Ontario, were therefore associated with lower accumulated vernalization, while the sites that received the most consistent year-to-year vernalization were coastal midAtlantic sites at Warsaw, Virginia and Queenstown and Clarksville, Maryland (also shown by consistently low *vrn1* effect at Warsaw in Figure 6). Gradient descent of the combined model converged at a cumulative vernalization breakpoint of 83.4, above which no additional effect of increasing vernalization on heading date was estimated.

**Figure 3.**
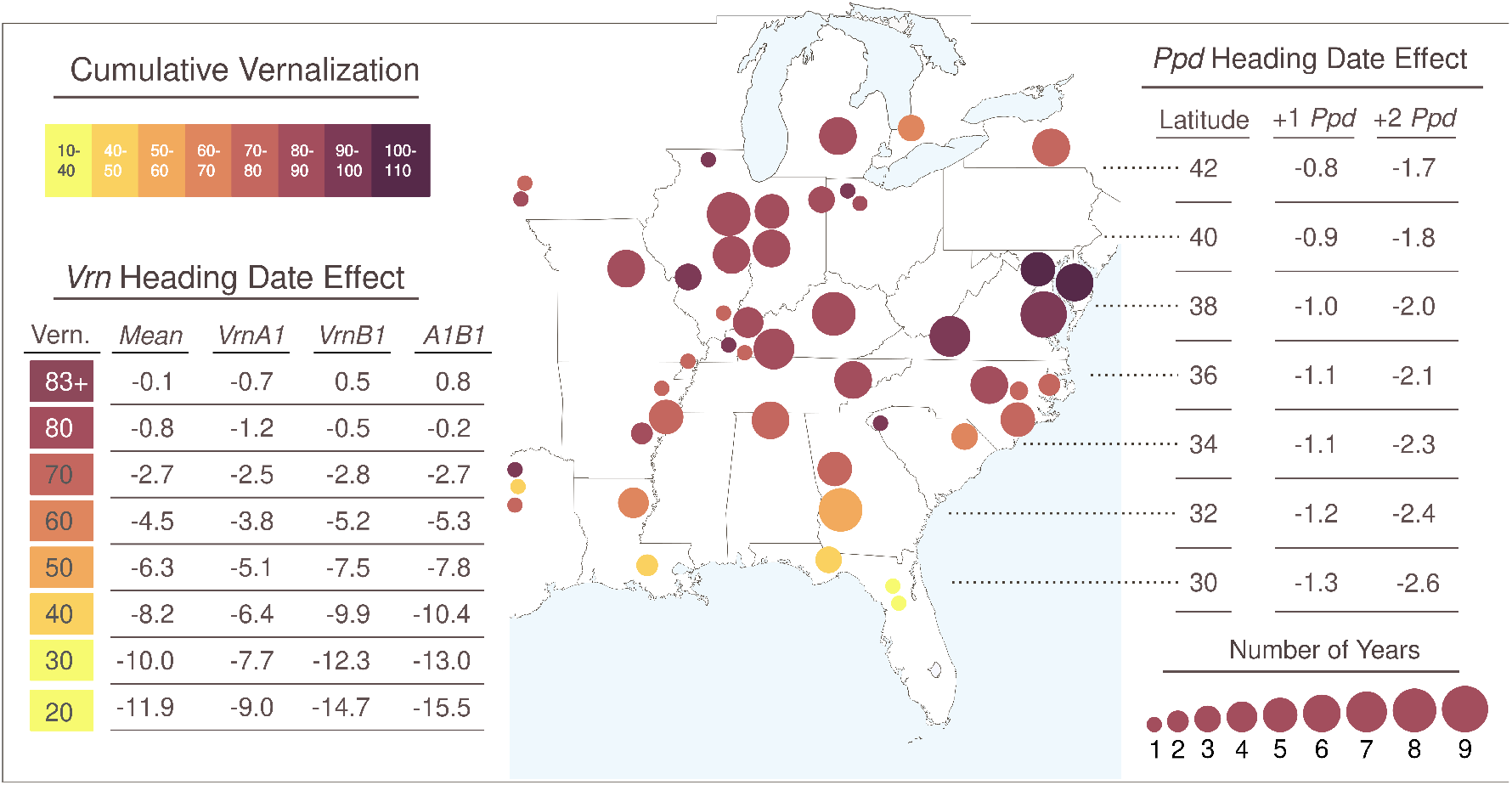
Major HD allele effects in tested sites. Center - map of locations in data set. Size of circles marking locations corresponds with the number of observations associated with each location across 8 years of data, and the color is associated with the average accumulated vernalization across all years. Left - estimated effects on heading date (days) of weak winter alleles at either *VrnA1* or *VrnB1* both from the combined model given different quantities of accumulated vernalization. Right - estimated effects on heading date (days) of one or two additional *Ppd*1 photoperiod insensitivity alleles given different latitudes.

In the combined allele effect model, both vernalization class and photoperiod insensitive allele dosage were highly significant (*p ≈* 0 in Wald test). Predicted effects from the combined model under different environmental conditions show that the heading date effect of any of the weak vernalization classes at the *vrn1* loci is strongly determined by the amount of accumulated vernalization during the winter (*p ≈* 0, Fig. 3). An additional mean effect of having any of the weak vernalization alleles was calculated from a weighted average of the individual weak class effects (weak *vrnA1*, weak *vrnB1, vrnB1* and *vrnB1*). In some years in southern environments where accumulated vernalization ranges from 20-30, the predicted effect of vernalization insensitivity on heading date was large – from 9 to 16 days earlier heading. This conforms with breeder observations from low-vernalization years at Baton Rouge, LA and Citra, FL, where the heading dates of strong and weak vernalization lines differ by approximately two weeks. As accumulated vernalization increases towards values common in the South and occasionally encountered in the northern nursery locations (40-60), the effect on heading date decreases in magnitude but remains large, in the range of 4 to 10 days. Fully vernalized environments common in Midwestern locations and nearly universal in the coastal Mid-Atlantic (sites in Maryland and Virginia) were associated with predicted effects of approximately zero. In most years at these locations, there was almost no difference in heading date between the strong and weak vernalization classes.

Photoperiod effects were comparatively consistent across latitudes; a single additional homozygous photoperiod allele was associated with a 0.8 day decrease in heading date at the northernmost latitude of 42 degrees, and a 1.3 day decrease in heading date at the southernmost latitude of 30 degrees (*p* = 0.02 for interaction effect, Fig. 3). This is consistent with previous estimates of *PpdD1* effects by Guedira et al. (2016) and DeWitt et al. (2021). Differences in day length at a given reproductive stage, which could alter effect of additional photoperiod insensitivity alleles from year to year within a single site at the same latitude as mean heading dates varied at that site, were not modeled.

### Heading Date Differences Create Yield Differences

Tested markers had substantial effects on heading dates. If differences in heading dates generate variation in yield, then these alleles will also be drivers of yield differences. To investigate evidence for the importance of heading date differences in driving GxE effects on wheat yield in this panel, correlations between heading date and yield within each of 445 site-years were calculated and displayed (Figure 4). The most common relationship was a slightly negative correlation (*e*.*g*. Kinston, NC 2018), but in many environments there was either a stronger negative correlation (*e*.*g*. Winnsboro, LA 2008), or even a positive one (*e*.*g*. Georgetown, MD 2008). While the overall relationship between heading date and yield is slightly negative, this obscures the underlying pattern of stronger relationships that vary by location and year, where heading date is often a major driver of entry-mean yield differences.

**Figure 4.**
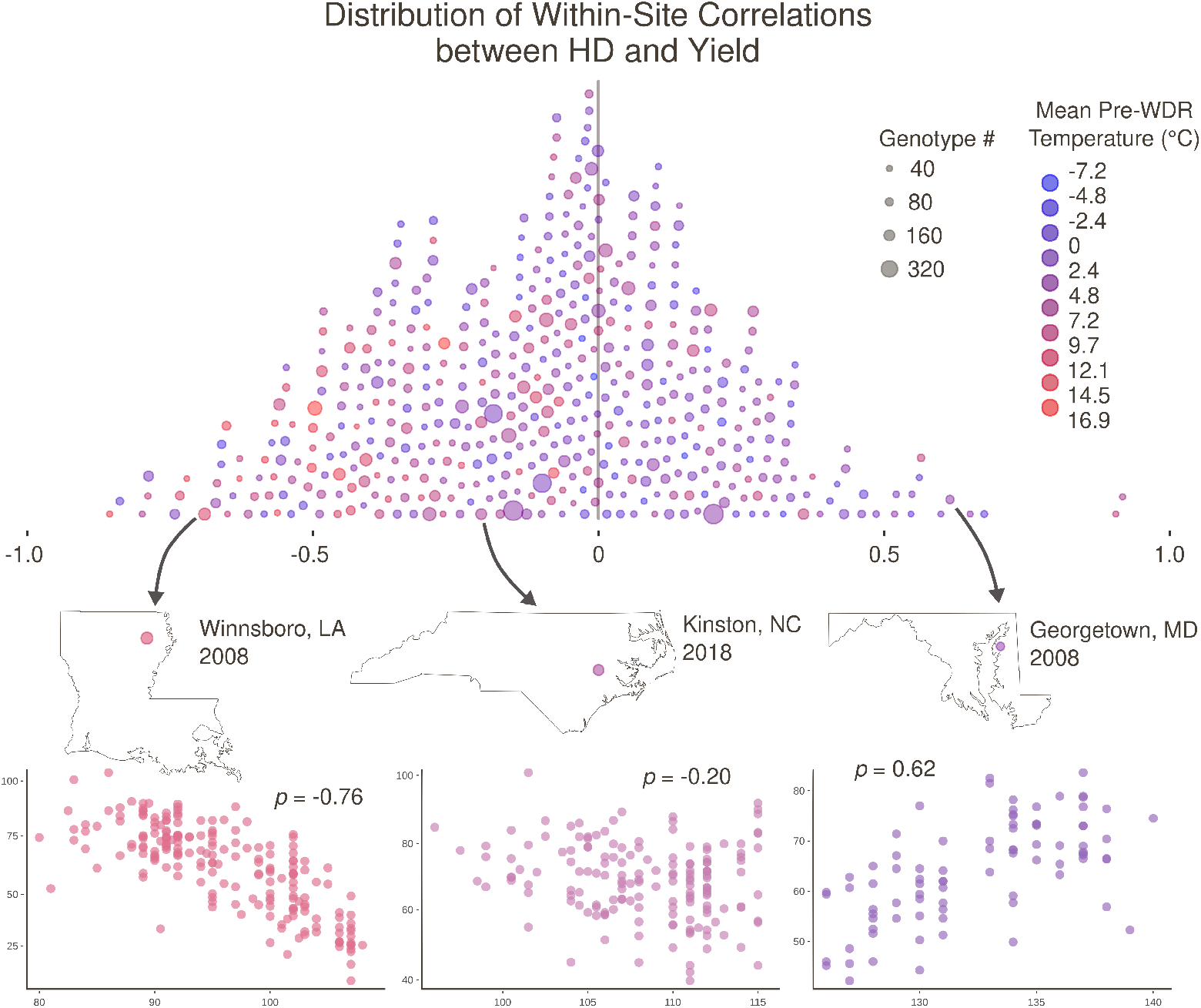
Distribution of site-year specific correlations between heading date and yield. For each site-year, correlations were calculated between genotype values for heading date and yield and plotted. Circles are scaled relative to the number of lines included in the site-year, and are colored by mean winter temperature. The mean correlation is slightly negative, but environments with both strongly positive and negative correlations between HD and yield were observed.

We evaluated the importance of different heading date-specific photoperiod-dependent variables on grain yield variation using 1,000 iterations of a Bayesian lasso model that tested for the impact of those variables on yield. These photoperiod-dependent variables that differed for each unique heading date allowed for variation for effects *within* each site, instead of assigning a single environmental value to all entry-means at a given site-year. A set of 19 photoperiod-dependent variables that were almost always included in the model were identified (coefficient set to 0 in ≤ 1% of iterations). These variables included: average and maximum temperature during floral development, flowering, and grain fill, minimum temperature during floral development and grain fill, average precipitation and maximum precipitation during floral development and flowering, mean soil moisture during floral development and flowering, freeze damage during floral development and flowering, received shortwave radiation during flowering and grainfill, and maximum windspeed during floral development.

Selected variables were fit as mediating variables between relative heading date and yield across all locations in a single mediation analysis model to understand how photoperiod-dependent variables mediated the within-environment yield effects of heading date (Figure 5). Later heading increased the temperatures plants experienced during all three developmental stages (vegetative, flowering, and grain fill), for average temperature, minimum temperature (equivalent to greater night-time temperatures), and maximum temeprature (equivalent to greater day-time temperatures). The increase in average temperature during vegetative development was associated with a decrease in relative yield, while increases in minimum and maximum temperature relative to average temperature were associated with increases. Later heading was associated with small average decreases in soil moisture, decreases that are associated with increased yield when experienced during vegetative growth but decreased yield during flowering and grain fill. Increases in solar radiation during flowering and grain fill were associated with later heading, which increased yield during flowering but decreased yield during grain fill. Delayed heading decreased yield losses due to freeze damage by decreasing mean freeze damage during vegetative growth and flowering, both of which were associated with negative effects on yield.

**Figure 5.**
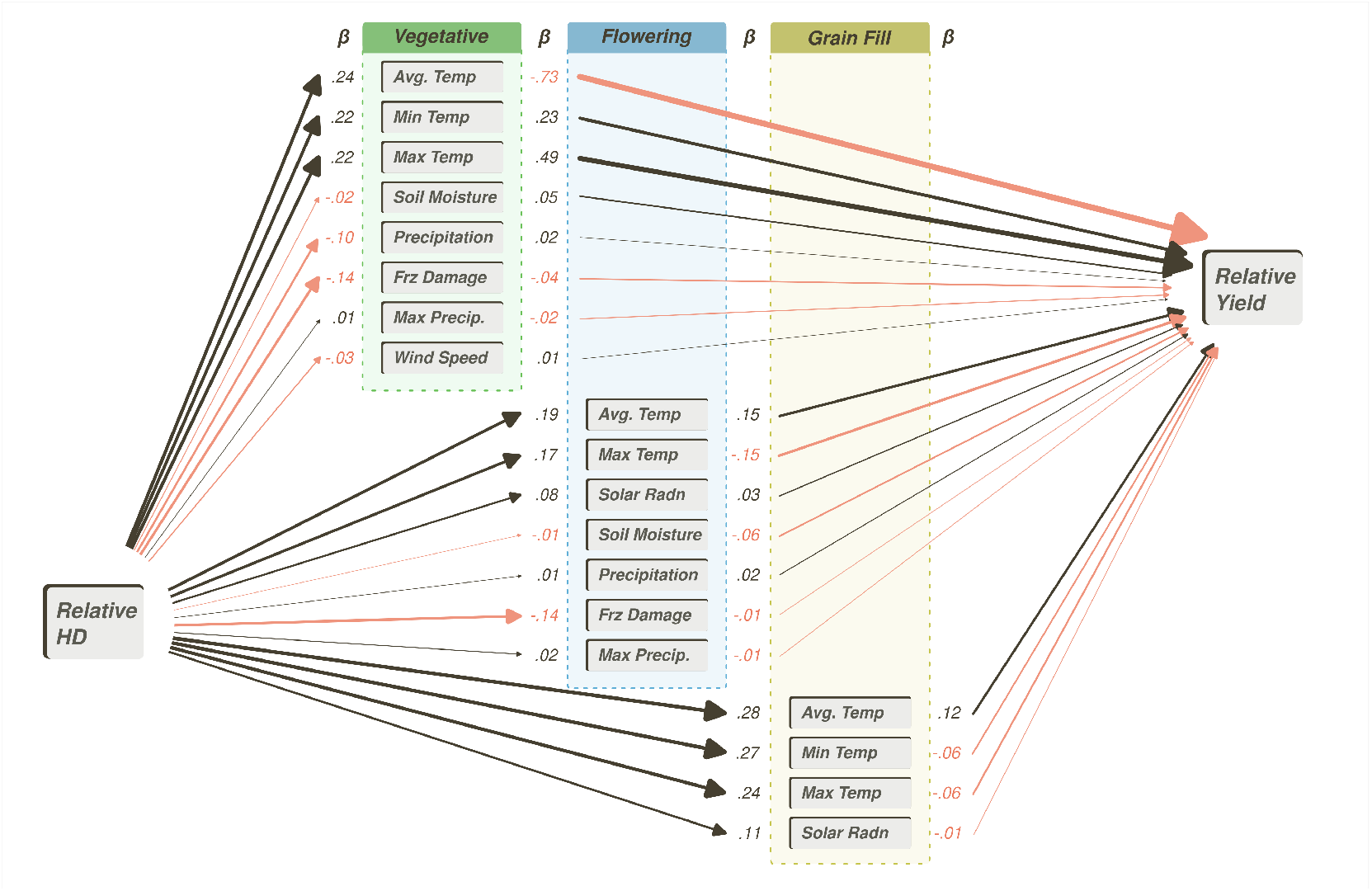
Estimated effects of heading date differences on yield. Effects of HD on yield in a location-specific context were estimated as both the direct effect of relative heading date on yield and on the effect mediated by stress variables. Not shown are effects of principal components of the **H** matrix on yield.

**Figure 6.**
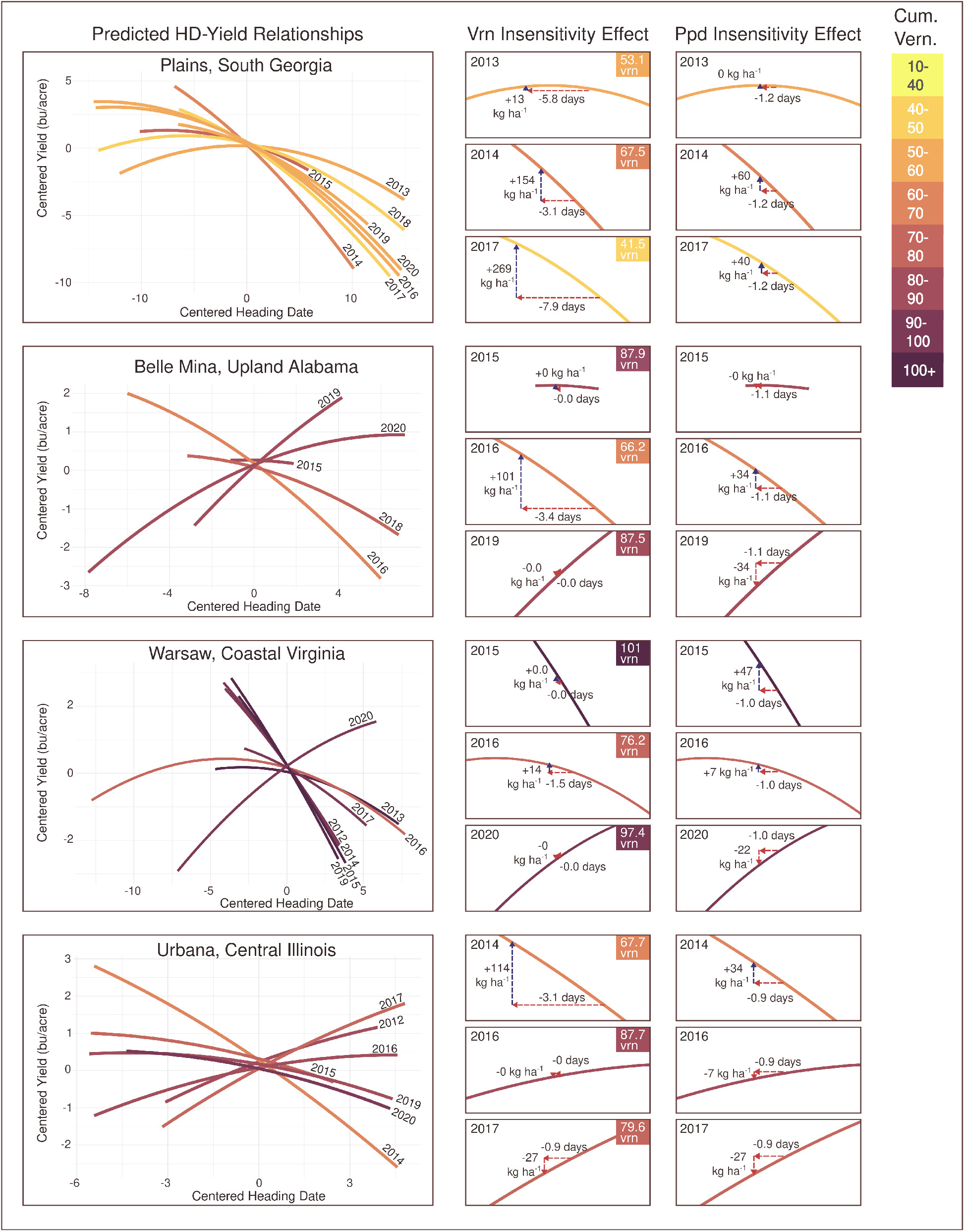
Predicted HD allele effects in select environments. Estimated heading date and yield relationships for selected environments as predicted by phenologically dependent variables. Combined effects of vernalization insensitivity and additional photoperiod insensitive allele are modeled as a function of accumulated vernalization and latitude, respectively, and the within-site relationships are used to estimate a yield effect from the heading date effect for each site-year.

**Figure 7.**
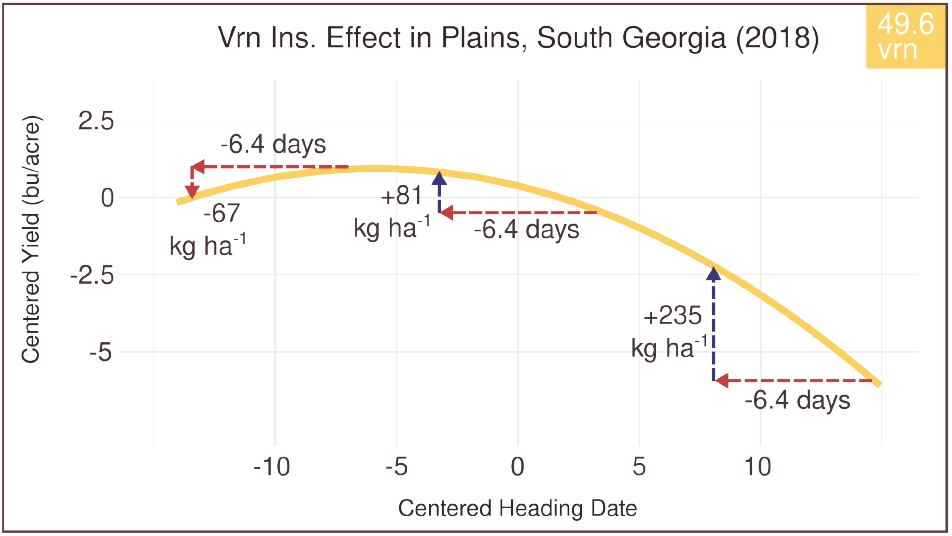
Yield effects of weak vernalization alleles in different phenotypic contexts. Relationship between heading date and yield in an example environment with low vernalization (here, 49.5 accumulated vernalization). Insufficient vernalization of vernalization-sensitive lines is predicted to have different yield effects depending on the expression of other heading date alleles.

### Mediated Yield Effects of Heading Date Genes

Estimates from a model that fit relationships between relative (centered) heading date and yield within each location were used to fit curves demonstrating the predicted relationship between heading date and yield in each site-year as a function of the photoperiod-dependent variables associated with each observed heading date value at that site-year (Fig. 6). A range of relationships between heading date and yield are predicted for each location – some strongly positive, some strongly negative, and a range of intermediate relationships, including some where optimal heading dates are at values closer to zero. In general, more southern locations tended to have less accumulated vernalization, and tended towards more frequent negative relationships between heading date and yield (later-maturity lines yielding less).

The effect of vernalization insensitivity and photoperiod insensitivity on yield were predicted for each of these locations given the associated accumulated vernalization and latitude, respectively. For each predicted heading date difference relative to the mean heading date, a predicted yield difference was estimated based on the predicted within-site-year relationships (Fig. 6). In general, the yield effect of a weak vernalization allele tended to be either close to zero, or larger and positive, with few cases of strong negative effects on yield. This is because the same locations where the vernalization effect was large (site-years with lower accumulated vernalization) tended to be those same site-years in which later lines were predicted to perform better. This was not the case for the photoperiod alleles, where the overall effect was modeled as consistent within each site-year. Because of this, the photoperiod alleles produced yield effects that were smaller than the larger yield effects of the *vrn1* alleles, but larger in the near-zero vernalization effect site-years where full vernalization was achieved. Earlier heading due to multiple photoperiod insensitivity alleles was associated with both positive, negative, and nearzero yield effects depending on the predicted relationship between heading date and yield at that site-year. Practically, in the assayed environments, vernalization insensitivity alleles generally promoted greater yield stability, while photoperiod insensitivity alleles generally promoted greater yield instability.

## Discussion

### Vernalization and Photoperiod Insensitivity Effects through Heading Date

Differences in heading date often help drive relative differences in genotype performance between environments (as an example for these specific alleles, Addison et al. (2016)). In our trial data, we observe a distribution of correlations between heading date and grain yield in 445 different site-years that supports the anecdotal observation that this relationship varies, is often strong, and is more likely to be negative instead of positive in more southern, warmer climates (Fig. 4). Heading date differences, representing overall differences in maturity between lines, help determine differences in yield ranking between individuals. The strength of this relationship and its variability suggests that better modeling of this relationship and its genetic and environmental basis may improve yield predictions.

Both *vrn1* and *Ppd1* alleles have been tools used by Eastern wheat breeders to adjust their lines’ heading date habits and average yield (Addison et al., 2016; Mason et al., 2018; Sarinelli et al., 2019). In the South, the use of weak winter alleles of *vrn1* has been critical to prevent delayed heading in years where very little vernalization is accumulated due to warm winters. Figure 6 demonstrates how these early-heading alleles can have large positive effects on yield in environments with strong negative relationships between heading date and yield. Earlier heading associated with *Ppd1* insensitivity is consistent from year to year, which leads to positive yield effects when the relationship between heading date and yield is negative and negative yield effects when the relationship between heading date and yield is positive. Earliness associated with the presence of more than one *Ppd1* insensitivity allele, while comparatively common in northern breeding nurseries, can therefore be detrimental in locations where early-flowering lines generally perform worse due to environmental factors like late spring freezes. Conversely, the positive effects of *vrn1* alleles tend to be larger in low-vernalization environments where, due to their generally warmer weather, earlier lines typically yield better. Environments where a large early heading effect could be detrimental are, in turn, more likely to be cooler environments associated with greater accumulated vernalization and small or near-zero earliness effects on heading date and therefore yield. In the model used in this analysis, multiple *Ppd1* insensitivity alleles may promote greater yield instability, increasing yield in some environments and decreasing it in others, while *vrn1* earliness is rarely detrimental and may promote greater yield stability. Despite this seemingly advantageous habit, weak *vrn1* alleles are much less frequent in northern breeding programs than the *Ppd1* insensitivity alleles. Baring additional tradeoffs not captured by this analysis, northern programs may consider greater use of weaker *vrn1* alleles to promote earlier heading in advantageous environments, especially as winters tend towards increasing temperatures and greater temperature variability. However, c aution should be exercised when using weak *vrn1* alleles in combination with *PpdD1a* or multiple *Ppd1* insensitive alleles, instead of a weaker *PpdA1a*.*1* or *PpdB1a* alleles alone.

### Heading Date Alleles as Environmentally Conditional Yield QTL

The effects of heading date alleles on yield are intuitive to breeders but not well captured by the additive genomic prediction models that now inform selection decisions. In practice, genomic selection training populations are often constituted of large collaborative trials which span a collection of environments much wider than any individual breeder’s target population of environments. Across the large number of years and locations in these well-developed training population data sets, the positive and negative yield effects of an increase or decrease in heading date cancel out, so that estimates of the overall additive effects of major heading date alleles on yield over the entire training population may be quite small. In that sense, these alleles may generate a very small portion of the overall true variation in across-location breeding values while generating a major portion of the true variation in location-specific and especially site-year-specific breeding values. The inability of traditional modeling approaches to adequately capture these site-year-specific effects results in part from the low correlation between the true site-year breeding values and the true multi-location breeding values.

An additional complication arising from the heading date to yield relationship is the epistasis it generates under the assumption of a lack of linearity between heading date and yield. Considering the single site-year of Plains 2018: the difference in slope at different values of centered heading date generate differences in yield effects for the same heading date allele effect, depending on the “starting position”of the genotype within which the allele substitution occurs. In this instance, the same decrease in heading date will generate different additive effects conditional on the context of the phenotypic expression of all other heading date alleles packaged in that genotype. If those other alleles plus a vernalization sensitive allele would place the line at the right part of the tail, further earliness would have large positive effects on the additive yield value. In the context of an allelic package that would otherwise generate an earlier phenotypic value in the context of that environment, the additive yield effect of vernalization insensitivity would instead generate a near-zero or negative yield effect. This epistasis in yield effect even under the assumption that the heading date alleles have purely additive effects on heading date generates additional complications in modeling heading date allele effects on yield (Mason et al., 2018).

The availability of large-scale weather data has enabled environment-specific genomic predictions as functions of weather and marker data (Heslot et al., 2014; Rogers and Holland, 2021). Some modeling approaches have been able to integrate modeling of developmental-stage variables to improve the performance of these models (Millet et al., 2019). Given the complicated interactions between heading date and yield, future work is required that integrates explicit modeling of these major genes and their interactions with weather variables alongside genome-wide marker-based predictions to improve prediction accuracy for yield. Explicitly accounting for major gene effects should improve the estimation of additive yield effects generated by those alleles in the target population of environments, and improve estimation of other allele effects by adjusting for the sometimes major within-trial yield effects of these major heading date alleles. Additional collection and centralization of data sets for which phenotype, genotype, and weather data are available will facilitate this model development.

### Implications for Breeding Programs

Marker data that assays *vrn1* and *Ppd1* alleles is now generally available to wheat breeders. For other genes, such as those conferring disease resistance or increased quality, the relationship between this data and breeding utility is straightforward – breeders can select on the allele with the desired effect. The complicated yield effects of these alleles observed in this study demonstrates why the use of heading date alleles has been more complicated, as their effect on both heading date itself and yield values that are affected by heading date change from environment to environment. Two general approaches may be suggested for making use of this data: characterization of lines’ *vrn1* and *Ppd1* multi-locus haplotypes to predict their general behavior in a specific subset of environments, and utilization of marker information to extend genomic prediction models to accommodate nonadditive effects generated by these genes.

As detailed in Figure 3, a single copy of any of the vernalization alleles is sufficient to induce a qualitative change in plant phenology, and the near-absence of wheat lines suggests the same is true for photoperiod alleles. Additional alleles may quantitatively increase or decrease heading date after the initial change, but the primary piece of information needed by a breeder to understand their line’s behavior is whether it has either of the vernalization alleles or any of the photoperiod insensitivity alleles. The very low rates of total photoperiod sensitivity suggest that the phenology of those lines is selected against in early breeding program stages. Common multi-location additive effects of additional photoperiod insensitivity alleles, however, will contribute much more to variation in heading date in northern environments where vernalization requirements are generally met and the overall variance in heading date is much lower; for Southern breeders then, the primary alleles of interest may be grouped to classify lines into vernalization sensitive vs insensitive based on the multi-location haplotype at the two *V rn*1 loci, while for more Northern breeders the number of photoperiod insensitivity alleles may be the primary concern. Additional photoperiod insensitivity alleles, however, also appear to have more potential to generate negative yield variability than vernalization insensitivity. Low-vernalization environments are generally associated with environmental conditions that favor earlier-heading lines, so that vernalization insensitivity tends to reduce variability for yield, while an early habit conferred by multiple photoperiod insensitivity alleles can punish lines in conditions where later lines are favored. Additional work may focus on longer-photoperiod, vernalization-insensitive lines that attempt to minimize yield variability with more variable weather conditions.

Incorporating weather data into genomic prediction models allows for the estimation of environment-specific effects that may improve performance of environment-specific predictions of additive merit. Reaction norm models have demonstrated utility in environment-specific prediction by fitting interactions, in various ways, between all markers and all season-long environmental variables (*e*.*g*. Jarquín et al. (2014), Heslot et al. (2014), Rogers and Holland (2021)). In this study, weather variables at all developmental stages are associated with yield variation, not just grain fill. General trends in average temperature appear more important than specific stress events like freeze damage, drought stress, and precipitation, although this may be a limitation of the data we have available. Given that heading date variation driven in major part by a small number of vernalization and photoperiod insensitivty alleles was important for generating environment-specific yield variation, explicit modeling of these interactions has the potential to improve reaction norm GxE models. These improvements might take the form of different weighting of markers known to be linked to these alleles, classification of allele data into qualitative sensitive/insensitive classes, and transformation of raw weather variable data into heading date-specific photoperiod-dependent variables (if heading date data is available).

## Conclusions

Wheat geneticists have uncovered major genes responsible for generating large effects on many phenotypes, including on maturity and heading date. Actually utilizing these genes in breeding has proven to be complicated. It is rarely the case that one allele is always favorable across genetic backgrounds and environments, and the relative importance of known alleles relates to their allele effects, which are hard to estimate and are contextual to the germplasm and testing set of environments. A necessary first step towards integrating information on major genes into breeding decisions involves actually estimating allele effects as they change from one environment to the next. Here we take initial steps to illustrate how alleles generating variation for wheat phenology can generate differential yield effects as a result of exposing plants to different environmental conditions. Population structure and lack of balance between panels limits the utility of point estimates of allele effects from this model. However, historic associations between lack of vernalization hours and a negative association between heading date and grain yield suggest that the use of *vrn* insensitivity alleles to achieve earlier heading may promote greater year-to-year yield stability. Variable selection points to a reduced number of developmental-stage specific weather variables – primarily temperature, precipitation, freeze damage, and solar radiation – that are actually relevant to generating environment-specific yield variation. Future models predicting genotype performance may treat yield as the outcome of a longitudinal process through which variation in plant development affects and is affected by environmental factors.

## Author’s contributions

ND aggregated historic data, conducted all statistical analyses, generated all figures, and wrote the initial draft. MG originated the concept of classifying genotypes by their vernalization and photoperiod class and performed preliminary analyses of data. GBG assisted with the conceptual framework of the manuscript and its analysis, wrote sections of the final draft, and provided extensive edits and feedback. JL and BW helped aggregate data sets for individual nurseries and provided feedback on analyses. CM assisted with mediation analyses and provided critical feedback on methods and their discussion. MAB, RB, SH, REM, EM, MMe, MMo, JPM, EO, JR, NS, CS, MS, RS, VT, and DAVS designed, planted, and conducted analyses of yield trials used in the manuscript, as well as providing edits and feedback on early drafts of the manuscript. KH and JHS generated all genotype data used in the manuscript. ZW and WS provided edits and wrote sections of the final manuscript. NB provided edits and gave feedback on the final draft of the manuscript.

## Acknowledgments

Partial funding was provided the Louisiana Soybean and Grain Research and Promotion Board, the North Carolina Small Grain Growers’ Association, and University of Georgia Research Foundation (UGARF) and Georgia Seed Development (GSD). This work was supported in part by the Agriculture and Food Research Initiative Competitive Grant 2022-68013-36439 (Wheat-CAP) from the USDA National Institute of Food and Agriculture. Mention of trade names or commercial products in this publication is solely for the purpose of providing specific information and does not imply recommendation or endorsement by the U.S. Department of Agriculture.

## Supplemental Data

**Table S1:**
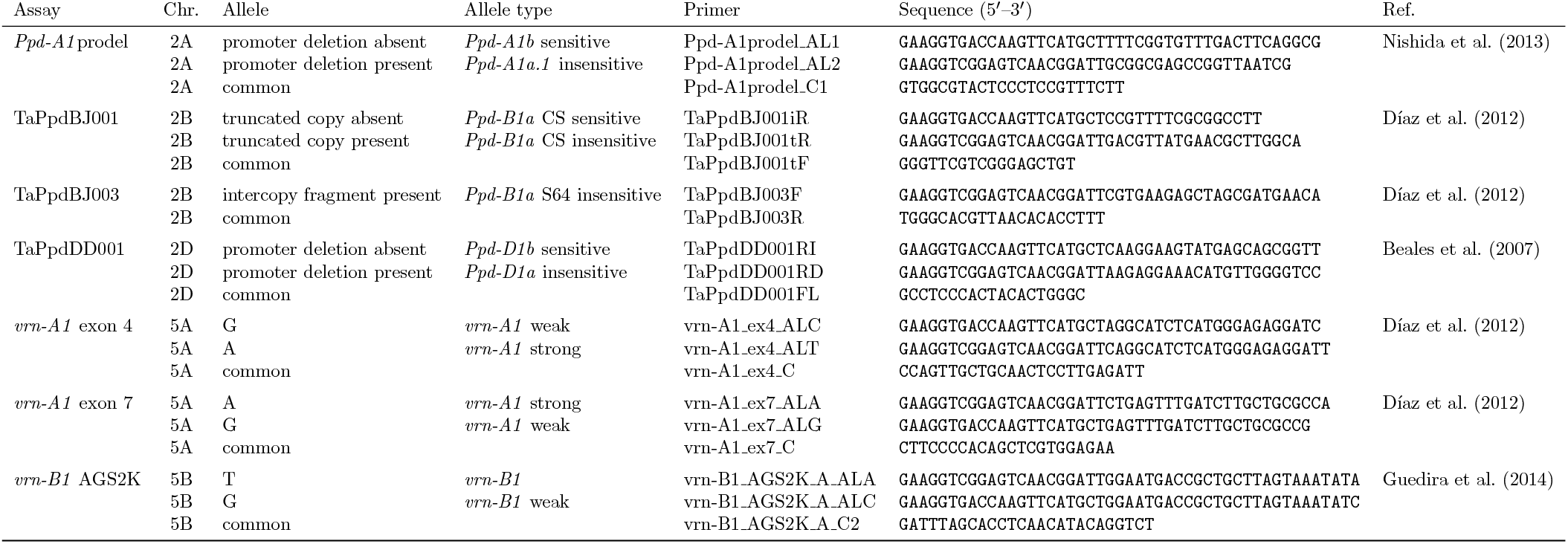
KASP Sequences for utilized markers. Sequences of Kompetititive allele specific primers for *vrn1* and *Ppd1* alleles, with polymorphism and effect noted, along with reference.

**Table S2:** Frequency of earlier flowering alleles at the Vrn1 and Ppd1 loci based on the KASP marker data set. Note that the total number of lines in this set is larger than the number used in subsequent analyses, as it includes lines that lacked genotypes at some of the loci but not others, and includes lines that do not have associated phenotypes.

|  | *vrnA1 weak* |  | *vrnB1 weak* |  | *PpdA1a.1* |  | *PpdB1a* |  | *PpdD1a* |  |
| --- | --- | --- | --- | --- | --- | --- | --- | --- | --- | --- |
|  | *prop.* | *n* | *prop.* | *n* | *prop.* | *n* | *prop.* | *n* | *prop.* | *n* |
| GAWN | 0.19 | 228 | 0.18 | 227 | 0.48 | 279 | 0.34 | 301 | 0.63 | 470 |
| SUNWheat | 0.26 | 360 | 0.29 | 357 | 0.41 | 368 | 0.46 | 351 | 0.67 | 368 |
| USSRWWN | 0.18 | 187 | 0.17 | 187 | 0.46 | 212 | 0.28 | 224 | 0.61 | 341 |
| Mason-Dixon | 0.17 | 228 | 0.03 | 227 | 0.52 | 294 | 0.22 | 301 | 0.74 | 349 |
| ABB Mid-Atlantic | 0.12 | 90 | 0.08 | 90 | 0.64 | 95 | 0.39 | 74 | 0.83 | 101 |
| Five State Pre | 0.01 | 157 | 0.00 | 157 | 0.71 | 170 | 0.17 | 182 | 0.51 | 216 |
| Five State Adv | 0.01 | 149 | 0.02 | 149 | 0.71 | 164 | 0.15 | 178 | 0.48 | 206 |
| UESRWWN | 0.07 | 172 | 0.02 | 172 | 0.59 | 197 | 0.22 | 214 | 0.56 | 343 |

## References

Christopher K. Addison, R. Esten Mason, Gina Brown-Guedira, Mohammed Guedira, Yuanfeng Hao, Randall G. Miller, Nithya Subramanian, Dennis N. Lozada, Andrea Acuna, Maria N. Arguello, Jerry W. Johnson, Amir M.H. Ibrahim, Russell Sutton, and Stephen A. Harrison. QTL and major genes influencing grain yield potential in soft red winter wheat adapted to the southern United States. Euphytica, 209(3):665–677, 6 2016. ISSN 15735060. doi: 10.1007/s10681-016-1650-1.

R. W. Allard and A. D. Bradshaw. Implications of Genotype-Environmental Interactions in Applied Plant Breeding 1. Crop Science, 4(5):503–508, 9 1964. ISSN 0011-183X. doi: 10.2135/crop-sci1964.0011183×000400050021x.

Richard G Allen, Luis S Pereira, Dirk Raes, Martin Smith, and others. Crop evapotranspiration-Guidelines for computing crop water requirements-FAO Irrigation and drainage paper 56. Fao, Rome, 300(9):D05109, 1998.

Rodrigo R. Amadeu, Catherine Cellon, James W. Olmstead, Antonio A. F. Garcia, Marcio F. R. Resende, and Patricio R. Muñoz. AGHmatrix: R Package to Construct Relationship Matrices for Autotetraploid and Diploid Species: A Blueberry Example. The Plant Genome, 9(3), 11 2016. ISSN 1940-3372. doi: 10.3835/plantgenome2016.01.0009.

James Beales, Adrian Turner, Simon Griffiths, John W. Snape, and David A. Laurie. A Pseudo-Response Regulator is misexpressed in the photoperiod insensitive Ppd-D1a mutant of wheat (Triticum aestivum L.). Theoretical and Applied Genetics, 115(5):721–733, 9 2007. ISSN 00405752. doi: 10.1007/s00122-007-0603-4.

SS Bhullar and CF Jenner. Differential Responses to High Temperatures of Starch and Nitrogen Accumulation in the Grain of Four Cultivars of Wheat. Functional Plant Biology, 12(4):363, 1985. ISSN 1445-4408. doi: 10.1071/PP9850363.

Richard E. Boyles, David S. Marshall, and Harold E. Bockelman. Yield Data from the Uniform Southern Soft Red Winter Wheat Nursery Emphasize Importance of Selection Location and Environment for Cultivar Development. Crop Science, 59(5):1887, 2019. doi: 10.2135/cropsci2018.11.0685.

Claire Dandine-Roulland and Hervé Perdry. Genome-Wide Data Manipulation, Association Analysis and Heritability Estimates in R with Gaston 1.5. In 46th European Mathematical Genetics Meeting (EMGM), page 6, Cagliari, Italy, 4 2018. Human Heredity. doi: 10.1159/000488519.

Noah DeWitt, Mohammed Guedira, Edwin Lauer, J. Paul Murphy, David Marshall, Mohamed Mergoum, Jerry Johnson, James B. Holland, and Gina Brown-Guedira. Characterizing the oligogenic architecture of plant growth phenotypes informs genomic selection approaches in a common wheat population. BMC Genomics, 22(1), 12 2021. ISSN 14712164. doi: 10.1186/s12864-021-07574-6.

Noah DeWitt, Jeanette Lyerly, Mohammed Guedira, James B Holland, J Paul Murphy, Brian P Ward, Richard E Boyles, Mohamed Mergoum, Md Ali Babar, Ehsan Shakiba, Russel Sutton, Amir Ibrahim, Vijay Tiwari, Nicholas Santantonio, David A Van Sanford, Kimberly Howell, Jared H Smith, Stephen A Harrison, and Gina Brown-Guedira. Bearded or smooth? Awns improve yield when wheat experiences heat stress during grain fill in the southeastern United States. Journal of Experimental Botany, 74(21):6749–6759, 11 2023. ISSN 0022-0957. doi: 10.1093/jxb/erad318.

Aurora Díaz, Meluleki Zikhali, Adrian S. Turner, Peter Isaac, and David A. Laurie. Copy number variation affecting the photoperiod-B1 and vernalization-A1 genes is associated with altered flowering time in wheat (Triticum aestivum). PLoS ONE, 7(3), 3 2012. ISSN 19326203. doi: 10.1371/journal.pone.0033234.

Dylan Beaudette, Jay Skovlin, Stephen Roecke, and Andrew Brown. soilDB: Soil Database Interface, 2021. URL https://CRAN.R-project.org/package=soilDB.

R. G. Flood and G. M. Halloran. The Nature and Duration of Gene Action for Vernalization Response in Wheat. Annals of Botany, 53(3), 3 1984. ISSN 1095-8290. doi: 10.1093/oxfordjournals.aob.a086700.

Daolin Fu, Péter Szűcs, Liuling Yan, Marcelo Helguera, Jeffrey S. Skinner, Jarislav Von Zitzewitz, Patrick M. Hayes, and Jorge Dubcovsky. Large deletions within the first intron in VRN-1 are associated with spring growth habit in barley and wheat. Molecular Genetics and Genomics, 273(1):54–65, 3 2005. ISSN 16174615. doi: 10.1007/s00438-004-1095-4. URL http://www.ncbi.nlm.nih.gov/pubmed/15690172.

Adam Gauley and Scott A. Boden. Genetic pathways controlling inflorescence architecture and development in wheat and barley. Journal of Integrative Plant Biology, 61(3):296–309, 3 2019. ISSN 1672-9072. doi: 10.1111/jipb.12732. URL https://onlinelibrary.wiley.com/doi/abs/10.1111/jipb.12

Robert B. Gramacy and Ester Pantaleo. Shrinkage regression for multivariate inference with missing data, and an application to portfolio balancing. Bayesian Analysis, 5(2):237–262, 7 2010.

Mohammed Guedira and Gary M. Paulsen. Accumulation of starch in wheat grain under different shoot/root temperatures during maturation. Functional Plant Biology, 29(4):495, 2002. ISSN 1445-4408. doi: 10.1071/PP01006.

Mohammed Guedira, Peter Maloney, Mai Xiong, Stine Petersen, J. Paul Murphy, David Marshall, Jerry Johnson, Steve Harrison, and Gina Brown-Guedira. Vernalization Duration Requirement in Soft Winter Wheat is Associated with Variation at the ¡i¿VRN-B1¡/i¿ Locus. Crop Science, 54(5):1960–1971, 9 2014. ISSN 0011-183X. doi: 10.2135/cropsci2013.12.0833.

Mohammed Guedira, Mai Xiong, Yuan Feng Hao, Jerry Johnson, Steve Harrison, David Marshall, and Gina Brown-Guedira. Heading date QTL in winter wheat (Triticum aestivum L.) coincide with major developmental genes VERNALIZATION1 and PHOTOPE-RIOD1. PLoS ONE, 11(5), 5 2016. ISSN 19326203. doi: 10.1371/journal.pone.0154242.

JS Hawker and CF Jenner. High Temperature Affects the Activity of Enzymes in the Committed Pathway of Starch Synthesis in Developing Wheat Endosperm. Functional Plant Biology, 20(2):197, 1993. ISSN 1445-4408. doi: 10.1071/PP9930197.

Nicolas Heslot, Deniz Akdemir, Mark E. Sorrells, and Jean Luc Jannink. Integrating environmental covariates and crop modeling into the genomic selection framework to predict genotype by environment interactions. Theoretical and Applied Genetics, 127(2):463–480, 2014. ISSN 00405752. doi: 10.1007/s00122-013-2231-5.

Diego Jarquín, José Crossa Xavier Lacaze, Philippe Du Cheyron, Joëlle Daucourt, Josiane Lorgeou, François Piraux, Laurent Guerreiro, Paulino Pérez, Mario Calus, Juan Burgueño, and Gustavo de los Campos. A reaction norm model for genomic selection using high-dimensional genomic and environmental data. Theoretical and Applied Genetics, 127(3):595–607, 3 2014. ISSN 0040-5752. doi: 10.1007/s00122-013-2243-1.

Chengzhi Jiao, Xiaoming Xie, Chenyang Hao, Liyang Chen, Yuxin Xie, Vanika Garg, Li Zhao, Zihao Wang, Yuqi Zhang, Tian Li, Junjie Fu, Annapurna Chitiki-neni, Jian Hou, Hongxia Liu, Girish Dwivedi, Xu Liu, Jizeng Jia, Long Mao, Xiue Wang, Rudi Appels, Rajeev K. Varshney, Weilong Guo, and Xueyong Zhang. Pan-genome bridges wheat structural variations with habitat and breeding. Nature, 637(8045):384–393, 1 2025. ISSN 0028-0836. doi: 10.1038/s41586-024-08277-0.

Nestor Kippes, Mohammed Guedira, Lijuan Lin, Maria A. Alvarez, Gina L. Brown-Guedira, and Jorge Dubcovsky. Single nucleotide polymorphisms in a regulatory site of VRN-A1 first intron are associated with differences in vernalization requirement in winter wheat. Molecular Genetics and Genomics, 293(5):1231–1243, 10 2018. ISSN 16174623. doi: 10.1007/s00438-018-1455-0.

C. N. Law and A. J. Worland. Genetic analysis of some flowering time and adaptive traits in wheat. New Phytologist, 137(1):19–28, 9 1997. ISSN 0028-646X. doi: 10.1046/j.1469-8137.1997.00814.x.

C N Law, J Sutka, and A J Worland. A Genetic study of day-length response in wheat. Heredity, 41(2):185–191, 10 1978. ISSN 0018-067X. doi: 10.1038/hdy.1978.87.

R. C. Lewontin. The Adaptations of Populations to Varying Environments. Cold Spring Harbor Symposia on Quantitative Biology, 22(0):395–408, 1 1957. ISSN 0091-7451. doi: 10.1101/SQB.1957.022.01.037.

Genqiao Li, Ming Yu, Tilin Fang, Shuanghe Cao, Brett F. Carver, and Liuling Yan. Vernalization requirement duration in winter wheat is controlled by TaVRN-A1 at the protein level. Plant Journal, 76(5):742–753, 12 2013. ISSN 09607412. doi: 10.1111/tpj.12326.

R. Esten Mason, Christopher K. Addison, Ali Babar, Andrea Acuna, Dennis Lozada, Nithya Subramanian, Maria Nelly Arguello, Randall G. Miller, Gina Brown-Guedira, Mohammed Guedira, and Jerry Johnson. Diagnostic Markers for Vernalization and Photoperiod Loci Improve Genomic Selection for Grain Yield and Spectral Reflectance in Wheat. Crop Science, 58(1): 242–252, 1 2018. ISSN 0011-183X. doi: 10.2135/crop-sci2017.06.0348.

Emilie J. Millet, Willem Kruijer, Aude Coupel-Ledru, Santiago Alvarez Prado, Llorenç Cabrera-Bosquet, Sébastien Lacube, Alain Charcosset, Claude Welcker, Fred van Eeuwijk, and François Tardieu. Genomic prediction of maize yield across European environmental conditions, 6 2019. ISSN 15461718.

Hidetaka Nishida, Tetsuya Yoshida, Kohei Kawakami, Masaya Fujita, Bo Long, Yukari Akashi, David A. Laurie, and Kenji Kato. Structural variation in the 5 upstream region of photoperiod-insensitive alleles Ppd-A1a and Ppd-B1a identified in hexaploid wheat (Triticum aestivum L.), and their effect on heading time. Molecular Breeding, 31(1):27–37, 7 2013. ISSN 13803743. doi: 10.1007/s11032-012-9765-0.

Gary M. Paulsen and Elmer G. Heyne. Grain Production of Winter Wheat after Spring Freeze Injury. Agronomy Journal, 75(4):705–707, 7 1983. ISSN 0002-1962. doi: 10.2134/agronj1983.00021962007500040031x.

R Core Team. R: A Language and Environment for Statistical Computing, 2021.

Anna R Rogers and James B Holland. Environment-specific genomic prediction ability in maize using environmental covariates depends on environmental similarity to training data. G3 Genes—Genomes—Genetics, 12 2021. ISSN 2160-1836. doi: 10.1093/g3journal/jkab440.

J. Martin Sarinelli, J. Paul Murphy, Priyanka Tyagi, James B. Holland, Jerry W. Johnson, Mohamed Mergoum, Richard E. Mason, Ali Babar, Stephen Harrison, Russell Sutton, Carl A. Griffey, and Gina Brown-Guedira. Training population selection and use of fixed effects to optimize genomic predictions in a historical USA winter wheat panel. Theoretical and Applied Genetics, 132(4):1247–1261, 4 2019. ISSN 00405752. doi: 10.1007/s00122-019-03276-6.

I Sofield, LT Evans, MG Cook, and IF Wardlaw. Factors Influencing the Rate and Duration of Grain Filling in Wheat. Functional Plant Biology, 4(5):785–797, 10 1977. ISSN 1445-4408. doi: 10.1071/PP9770785.

Adam Sparks. nasapower: A NASA POWER Global Meteorology, Surface Solar Energy and Climatology Data Client for R. Journal of Open Source Software, 3(30):1035, 10 2018. ISSN 2475-9066. doi: 10.21105/joss.01035. URL https://doi.org/10.21105/joss.01035.

Robert Tibshirani. Regression Shrinkage and Selection Via the Lasso. Journal of the Royal Statistical Society Series B: Statistical Methodology, 58(1):267–288, 1 1996. ISSN 1369-7412. doi: 10.1111/j.2517-6161.1996.tb02080.x.

Adrian Turner, James Beales, Sebastien Faure, Roy P. Dunford, and David A. Laurie. The Pseudo-Response Regulator Ppd-H1 Provides Adaptation to Photoperiod in Barley. Science, 310(5750):1031–1034, 11 2005. ISSN 0036-8075. doi: 10.1126/science.1117619.

P. M. VanRaden. Efficient methods to compute genomic predictions. Journal of Dairy Science, 91(11):4414–4423, 2008. ISSN 15253198. doi: 10.3168/jds.2007-0980.

L. Yan, A. Loukoianov, G. Tranquilli, M. Helguera, T. Fahima, and J. Dubcovsky. Positional cloning of the wheat vernalization gene VRN1. Proceedings of the National Academy of Sciences, 100(10):6263–6268, 5 2003. ISSN 0027-8424. doi: 10.1073/pnas.0937399100.

L. Yan, M. Helguera, K. Kato, S. Fukuyama, J. Sherman, and J. Dubcovsky. Allelic variation at the VRN-1 promoter region in polyploid wheat. Theoretical and Applied Genetics, 109(8):1677–1686, 11 2004. ISSN 00405752. doi: 10.1007/s00122-004-1796-4.

L. Yan, D. Fu, C. Li, A. Blechl, G. Tranquilli, M. Bonafede, A. Sanchez, M. Valarik, S. Yasuda, and J. Dubcovsky. The wheat and barley vernalization gene VRN3 is an orthologue of FT. Proceedings of the National Academy of Sciences of the United States of America, 103(51):19581–19586, 12 2006. ISSN 00278424. doi: 10.1073/pnas.0607142103.

Yonggen Zhang and Marcel G Schaap. Weighted recalibration of the Rosetta pedotransfer model with improved estimates of hydraulic parameter distributions and summary statistics (Rosetta3). Journal of Hydrology, 547:39–53, 2017.

Bangyou Zheng, Karine Chenu, Alastair Doherty, and Scott Chapman. The APSIM-Wheat Module (7.5 R3008). Technical report, 2015. URL https://www.apsim.info/wp-content/uploads/2019/09/WheatDocumentation.pdf.

